# RNA localization to the mitotic spindle regulated by kinesin-1 and dynein is essential for early development of the sea urchin embryo

**DOI:** 10.1101/2022.08.16.504170

**Authors:** Carolyn M. Remsburg, Kalin D. Konrad, Jia L. Song

**Affiliations:** University of Delaware, Department of Biological Sciences, Newark, DE

**Keywords:** mitosis, RNA localization, kinesin-1, dynein, embryonic development

## Abstract

Mitosis is a fundamental and highly regulated process that acts to faithfully segregate chromosomes into two identical daughter cells. Transcript localization of genes involved in mitosis to the mitotic spindle may be an evolutionarily conserved mechanism to ensure that mitosis occurs in a timely manner. We identified many RNA transcripts that encode proteins involved in mitosis localized at the mitotic spindles in dividing sea urchin embryos and mammalian cells. Disruption of microtubule polymerization, kinesin-1, or dynein results in lack of spindle localization of these transcripts in the sea urchin embryo. Further, results indicate that the cytoplasmic polyadenylation element (CPE) within the 3’UTR of *Aurora B*, a recognition sequence of CPEB, is essential for RNA localization to the mitotic spindle. Blocking this sequence results in arrested development during early cleavage stages, suggesting that RNA localization to the mitotic spindle may be a regulatory mechanism of cell division that is important for early development.

## Introduction

Mitosis is the fundamental cellular process in which a cell divides to become two identical daughter cells following replication of its DNA (Mcintosh, 2016). This process involves the division of its duplicated DNA in karyokinesis and separation of the cytoplasm in cytokinesis (Mcintosh, 2016). The mitotic spindle is the organelle that drives the segregation of chromosomes (Gadde and Heald, 2004). The spindle is comprised primarily of tubulin monomers that heterodimerize (Petry, 2016). These monomers polymerize through the action of enzyme such as XMAP215, which is essential for the formation of mitotic spindles (Kronja *et al*., 2009). Actin also regulates mitosis by generating force within the dividing cell to orient the mitotic spindle, as well as separating chromosomes during anaphase (Anstrom, 1992; Kunda and Baum, 2009). Myosin II is the major motor protein that associates with actin and is indispensable for cytokinesis (Chaigne *et al*., 2016; Babkoff *et al*., 2021). Cofilin is an actin depolymerizer, whose inactivation is necessary for proper spindle orientation during mitosis, and is also responsible for importing actin into the nucleus (Pendleton *et al*., 2003; Kaji, Muramoto and Mizuno, 2008). Profilin is an actin-binding protein that is required for cytokinesis in chondrocytes, and increases actin export from the nucleus (Stüven, Hartmann and Görlich, 2003; Böttcher *et al*., 2009).

The segregation of chromosomes is highly dynamic and microtubule motors, such as kinesin-5, are required to slide anti-parallel microtubule fibers polewards (Cochran *et al*., 2005; Mann and Wadsworth, 2019). CENP-E, centromeric protein E, is a plus-ended kinesin motor protein that assists in orienting chromosomes properly along the metaphase plate (Craske and Welburn, 2020). Dynein is a microtubule minus-end directed motor protein that is known to regulate several aspects of mitosis, from centrosome separation to chromosome congression to spindle formation (Raaijmakers and Medema, 2014). Additionally, dynein and dynactin interact with NuMA to tether the astral microtubules to the cell cortex to orient the mitotic spindle (Hueschen *et al*., 2017; Okumura *et al*., 2018). NuMA also is essential for formation and maintenance of the spindle poles during mitosis (Zeng, 2000).

During early development, metazoan embryos undergo several rounds of rapid early cleavage divisions, where they cycle between mitosis (M) and synthesis (S) phases of the cell cycle, with minimal gap phases (Ikegami *et al*., 1994; Siefert, Clowdus and Sansam, 2015). Diverse cells accomplish mitosis in a relatively constant time frame of between 30 to 60 min, indicating exquisite regulation of mitosis to ensure a timely completion of this process (Araujo *et al*., 2016). Prolonged mitosis has been shown to result in cell death, cell arrest, or DNA damage (Rieder and Palazzo, 1992; Lanni and Jacks, 1998; Quignon *et al*., 2007; Uetake and Sluder, 2010; Orth *et al*., 2012). Thus, it is not surprising that mitosis is regulated by a plethora of mechanisms, from transcriptional regulation of cell cycle factors (Spellman *et al*., 1998; Whitfield *et al*., 2002) to post-translational regulation by phosphorylation and ubiquitination (Stegmeier *et al*., 2007; Dephoure *et al*., 2008; Lindqvist, Rodríguez-Bravo and Medema, 2009). In general, transcription is globally inhibited during mitosis (Martínez-Balbás *et al*., 1995); however, transcription occurs at the centromeric regions. Centromere transcription is essential for CENP-A nucleosome assembly and centromere formation and maintenance (Perea-Resa and Blower, 2018). Transcribed centromeric RNAs ensure correct CENP-C (RNA binding protein) levels and CENP-P (nucleosome) loading and accurate chromosome segregation. siRNAs and lncRNAs have also been found to be derived from centromeric regions and may play an important role in maintaining heterochromatin in the centromere domains (Hall *et al*., 2002; Volpe *et al*., 2002; Liu *et al*., 2015; Johnson *et al*., 2017; Perea-Resa and Blower, 2018).

As mitosis is under strict temporal control, it must be regulated in a rapid manner. As transcriptional regulation takes time, post-transcriptional and post-translational regulation play a key role during mitosis (Li and Zhang, 2017; Moura and Conde, 2019). Several critical steps in the cell cycle and specifically mitosis require post-translational regulation by kinases and phosphatases (Dephoure *et al*., 2008; Lindqvist, Rodríguez-Bravo and Medema, 2009; Combes *et al*., 2017; Gelens and Saurin, 2018; Moura and Conde, 2019). For example, entry into mitosis requires mitotic kinases Cdk1/Cyclin B1 and phosphatase Cdc25 (Boutros, Dozier and Ducommun, 2006; Lindqvist, Rodríguez-Bravo and Medema, 2009; Vigneron *et al*., 2018; Sun *et al*., 2019). Translation of cyclin B is required for sea urchin embryos to undergo mitosis (Chassé *et al*., 2016). Mitotic spindle elongation requires a perfect balance between kinase and phosphatase activities (Winey and Bloom, 2012; Nilsson, 2019). During metaphase, proper chromosome alignment is essential for progression through the spindle assembly checkpoint (SAC), which is passed partially through the inactivation of CDK1/Cyclin B and the activation of the anaphase-promoting complex/cyclosome (APC/C) via phosphorylation (Castro *et al*., 2005). Aurora B kinase also plays a role in regulating the transition from metaphase to anaphase, by phosphorylating several kinetochore components to reduce microtubule affinity and promote detachment from the kinetochore (Lens *et al*., 2003). In addition, phosphatases regulate APC/C activity by directly dephosphorylating Cdc20 and APC/C subunits and indirectly through phosphatase-mediated silencing of checkpoint signaling from the kinetochores to promote mitotic exit (Labit *et al*., 2012; Craney *et al*., 2016; Hein *et al*., 2017; Lee *et al*., 2017). Thus, the coordinated regulation of hundreds of proteins by dynamic phosphorylation during mitosis is a key mechanism to ensure the timely and precisely segregation of chromosomes.

While post-translational regulation of mitosis by kinases and phosphatases has been well-studied (Dephoure *et al*., 2008; Lindqvist, Rodríguez-Bravo and Medema, 2009; Combes *et al*., 2017; Moura and Conde, 2019), post-transcriptional regulation through localization of important RNA transcripts is less well understood. Several different proteins have been identified to control localization of RNAs to the mitotic spindle. Aurora B protein is recruited to the different areas of the kinetochores by phosphorylated histones where it has been found to associate with hundreds of mRNAs that are enriched on mitotic spindles, many of which encode cytoskeletal proteins and transcription factors (Jambhekar *et al*., 2014). The binding of Aurora B to mRNA is essential for its localization to centromeres, as well as its ability to phosphorylate its substrates, including Polo kinase, p53, securin, and APC/C, to initiate anaphase (Jambhekar *et al*., 2014). Staufen, an RNA binding protein, is thought to mediate localization of subpopulations of RNAs to the spindles during mitosis (Hassine *et al*., 2020). Additionally, Staufen regulates localization of *prospero* in *Drosophila* neuroblasts, ensuring asymmetric division and correct cell fate after mitosis (Broadus, Fuerstenberg and Doe, 1998). *Cyclin B* mRNA has previously been shown to have mitotic spindle localization in *Xenopus* and *Strongylocentrotus purpuratus*, as well as perinuclear localization in *Drosophila* embryos (Raff, Whitfield and Glover, 1990; Groisman *et al*., 2000; Yajima and Wessel, 2015). Interestingly, disruption of *Cyclin B* RNA localization results in defects in spindle architecture and ultimately abnormal cell division (Groisman *et al*., 2000). While many RNAs have been identified to localize to the mitotic spindle through biochemical assays (Blower *et al*., 2007; Sharp *et al*., 2011; Pascual *et al*., 2021), the idea that RNA localization is a more general mechanism for transcripts that encode proteins involved in mitosis has not been characterized in a developing embryo. In this study, we test the hypothesis that transcripts encoding proteins that regulate mitotic processes localize to the mitotic spindle, and that this localization is essential for early embryonic development.

We use the purple sea urchin embryo, *S. purpuratus,* to study RNA localization during mitosis. The sea urchin produces large and optically transparent blastomeres during cleavage stage which enables easy visualization of perturbation phenotypes (McClay, 2011). Using the sea urchin embryo and mammalian cells, we identified several transcripts that encode proteins involved in mitosis located on the mitotic spindle, indicating that this RNA localization is evolutionarily conserved. Transcripts that we examined were selected for their known roles in regulating the progression of mitosis. Further, this localization is dependent on microtubules and microtubule motor proteins kinesin-1 and dynein. Using reporter constructs, we also demonstrated that the 3’UTR of *Aurora B* is sufficient for spindle localization. Importantly, we identified a cytoplasmic polyadenylation element (CPE) sequence that is required for localization of *Aurora B* to the spindles and observed that blocking this CPE sequence results in developmental arrest. During cleavage stage development, the lack of gap phases during relatively rapid cell divisions may utilize spatial regulation of RNA transcripts of key players in mitosis to provide the cell another layer of control to an essential process. Our results reveal that RNA localization of *Aurora B* transcript to the spindles is a novel mechanism in ensuring proper development.

## Results

### RNA transcripts localize to the mitotic spindle

We examined the subcellular localization of the transcripts encoding select proteins involved in mitosis (Fig. 1A). The RNA transcripts of *Cyclin B*, *APC*, *Cdk*, *Aurora B*, *Polo kinase*, *CENP-E,* are localized between the dividing chromosomes of sea urchin 16 to 32 cell cleavage stage embryos (Fig. 1A). *NuMA* localizes to the astral microtubules and with the chromosomes.

**Figure 1:**
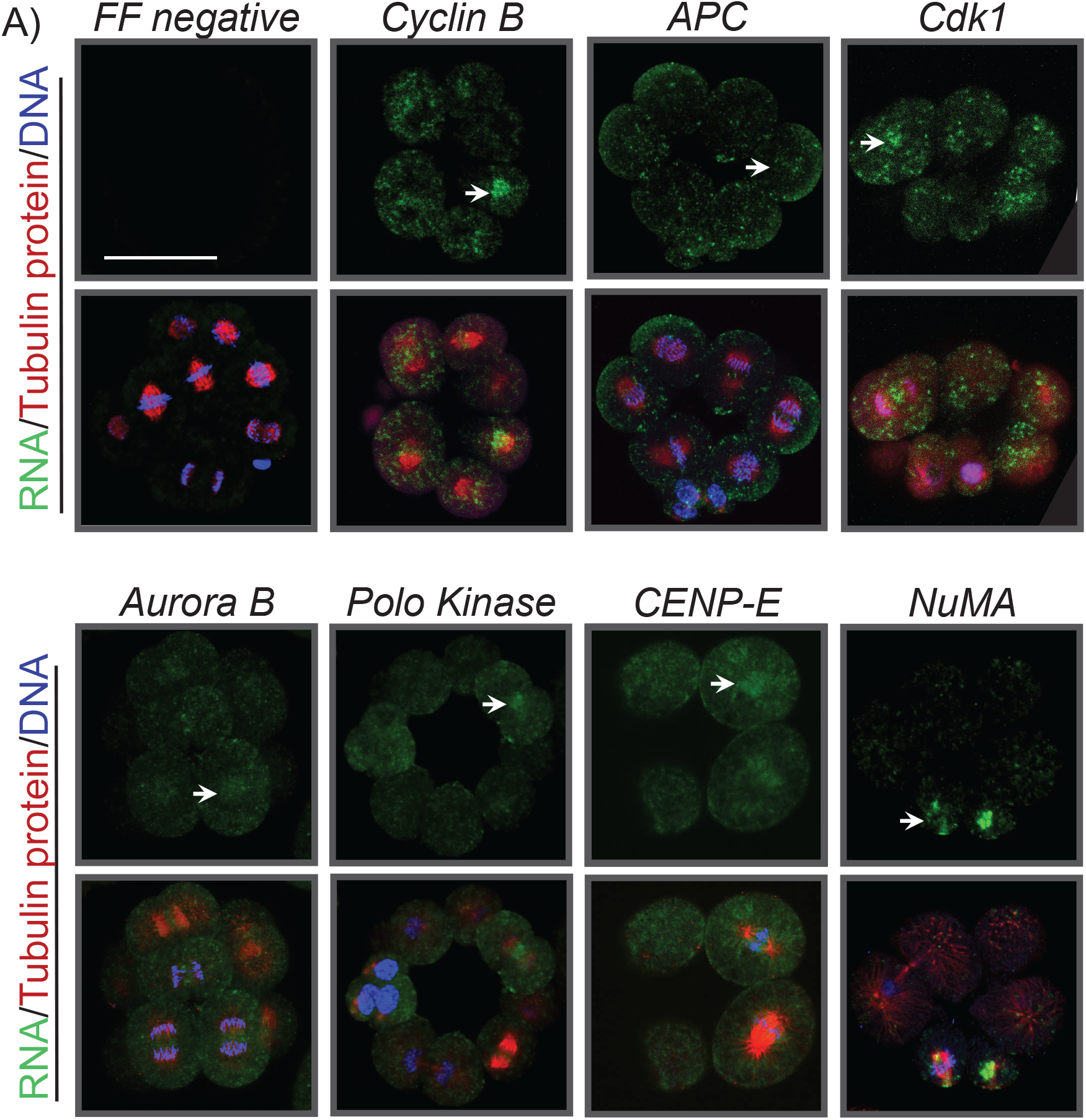
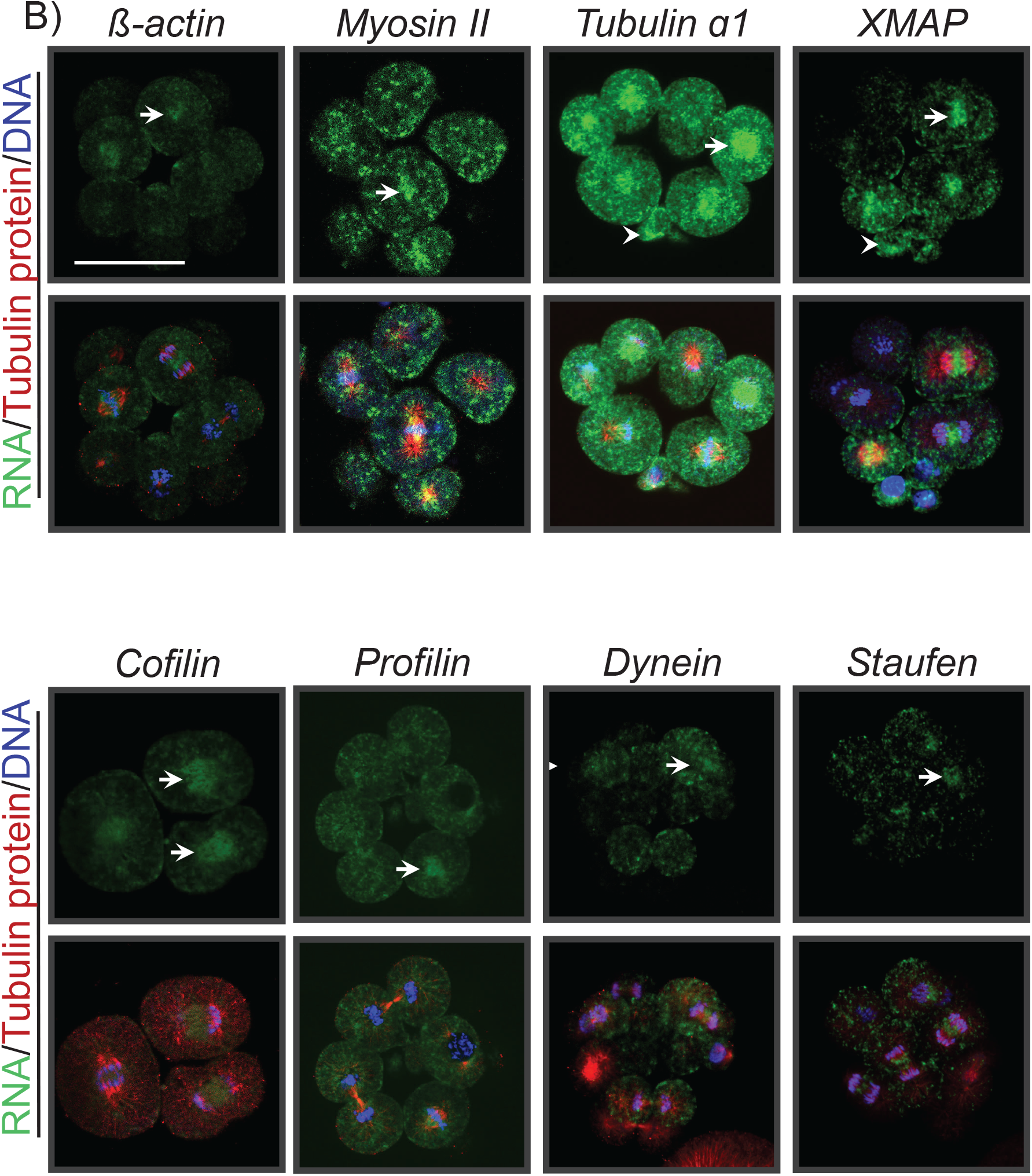
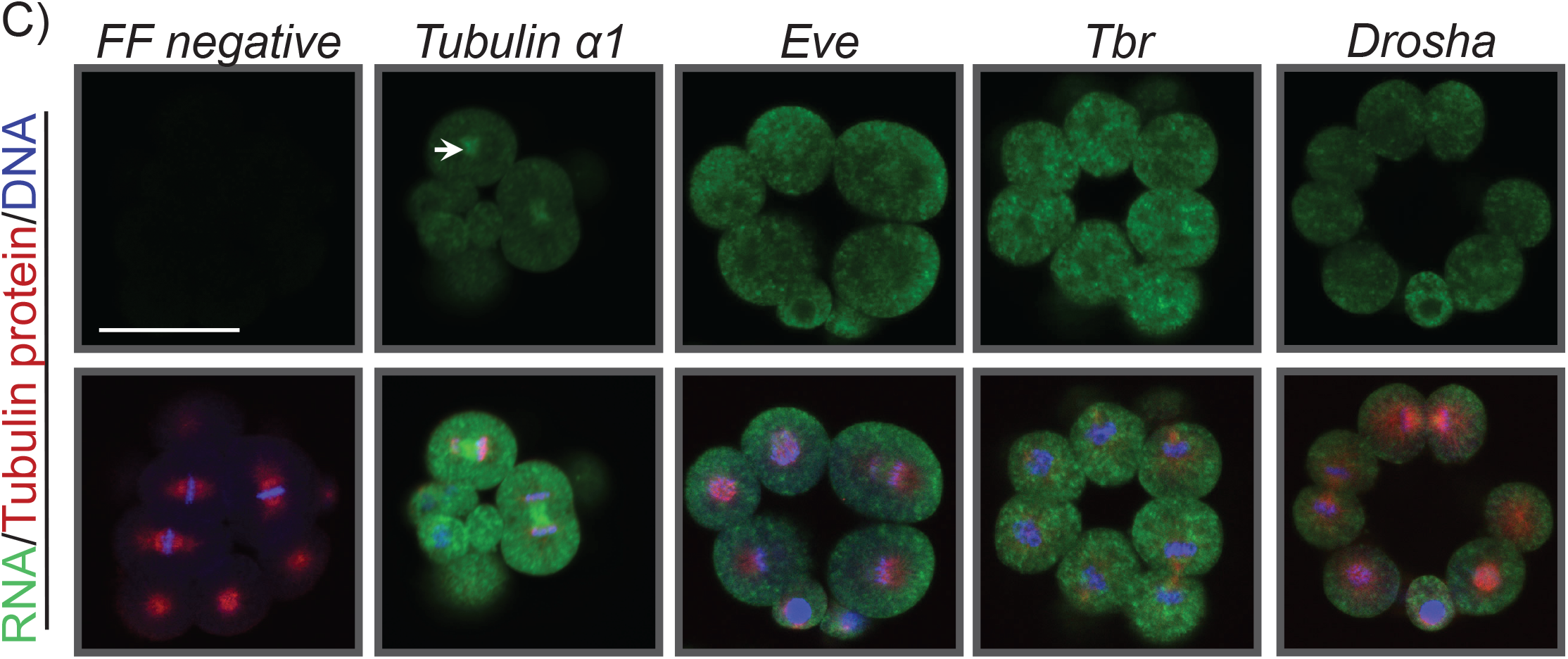
RNA transcripts that encode proteins that regulate mitosis localize to the mitotic spindle in developing embryos. (A) RNA transcripts that encode proteins that regulate mitosis localize to the mitotic spindle. (B) RNA transcripts that encode cytoskeletal proteins, motor proteins or transport RNA localize to the mitotic spindle. Embryos at the 16-32 cell stage were subjected to FISH, followed by immunolabeling with β-tubulin antibody, then counterstained with DAPI to detect DNA. Arrows indicate areas of RNA localization. Firefly (FF) is used as a negative control. Scale bar = 50 μm. (C) RNA transcripts that encode proteins that are not known to regulate mitosis and are expressed at the 16-32 cell stage do not localize to the mitotic spindle. Embryos were subjected to FISH using RNA probes, then immunolabeled with β-tubulin antibody, and counterstained with DAPI to detect DNA. *Tubulin α1* transcript is used as a positive control. Scale bar = 50 μm.

We also examined the subcellular localization of transcripts encoding cytoskeletal proteins, microtubule motor proteins, and regulators of other facets of mitosis (Fig. 1B). We observed that *β-actin*, *Myosin II*, *Tubulin α1*, *XMAP*, *Cofilin*, *Profilin*, *Dynein* and *Staufen* are localized between the dividing chromosomes of cleavage stage embryos (16 to 32 cells) undergoing mitosis (Fig. 1B). We also observe that the RNA transcripts of some of these genes (including *Aurora B, Polo kinase, β-actin, Tubulin α1, XMAP, Dynein,* and *Staufen*) localize to the perinuclear region in blastomeres in interphase (Fig. 1 and data not shown). These data indicate that the transcripts encoding proteins involved in mitosis are localized to the mitotic spindle.

To test the hypothesis that this localization of RNA transcripts to the mitotic spindle is selective and likely due to the function of these proteins in mitosis, we examined the subcellular localization of expressed transcripts that encode proteins of non-mitotic functions. Eve and Tbr are transcription factors that regulate endodermal and skeletal specification, respectively (Revilla-I-Domingo, Oliveri and Davidson, 2007; Peter and Davidson, 2010). Drosha is a dsRNA cleaving enzyme that processes microRNAs (Song *et al*., 2012). *Tbr* and *Drosha* are maternally present, while *Eve* is expressed by 6 hpf (Song and Wessel, 2007; Arshinoff *et al*., 2022). We observed that the RNA transcripts of *Eve, Tbr,* and *Drosha* are diffused throughout the cytoplasm of dividing cells (Fig. 1C). These data indicate that the localization of RNA transcripts to the mitotic spindle is not a general phenomenon that occurs with all RNA transcripts in the cell, but rather, a regulated process.

To test if this RNA localization is evolutionarily conserved in other organisms and cells, we examined subcellular localization of a select set of these transcripts in pig epithelial kidney cells (LLC-PK1) (Hull, Cherry and Weaver, 1976). We observed *Aurora B*, *Polo kinase 1* and *Staufen1* to localize between dividing nuclei in LLC-PK1 cells (Fig. 2). In interphase cells, *Staufen1* localizes to the perinuclear region (arrowheads in Fig. 2). These data indicate that the localization of RNA transcripts encoding proteins involved in mitosis is conserved from sea urchins to mammals.

**Figure 2:**
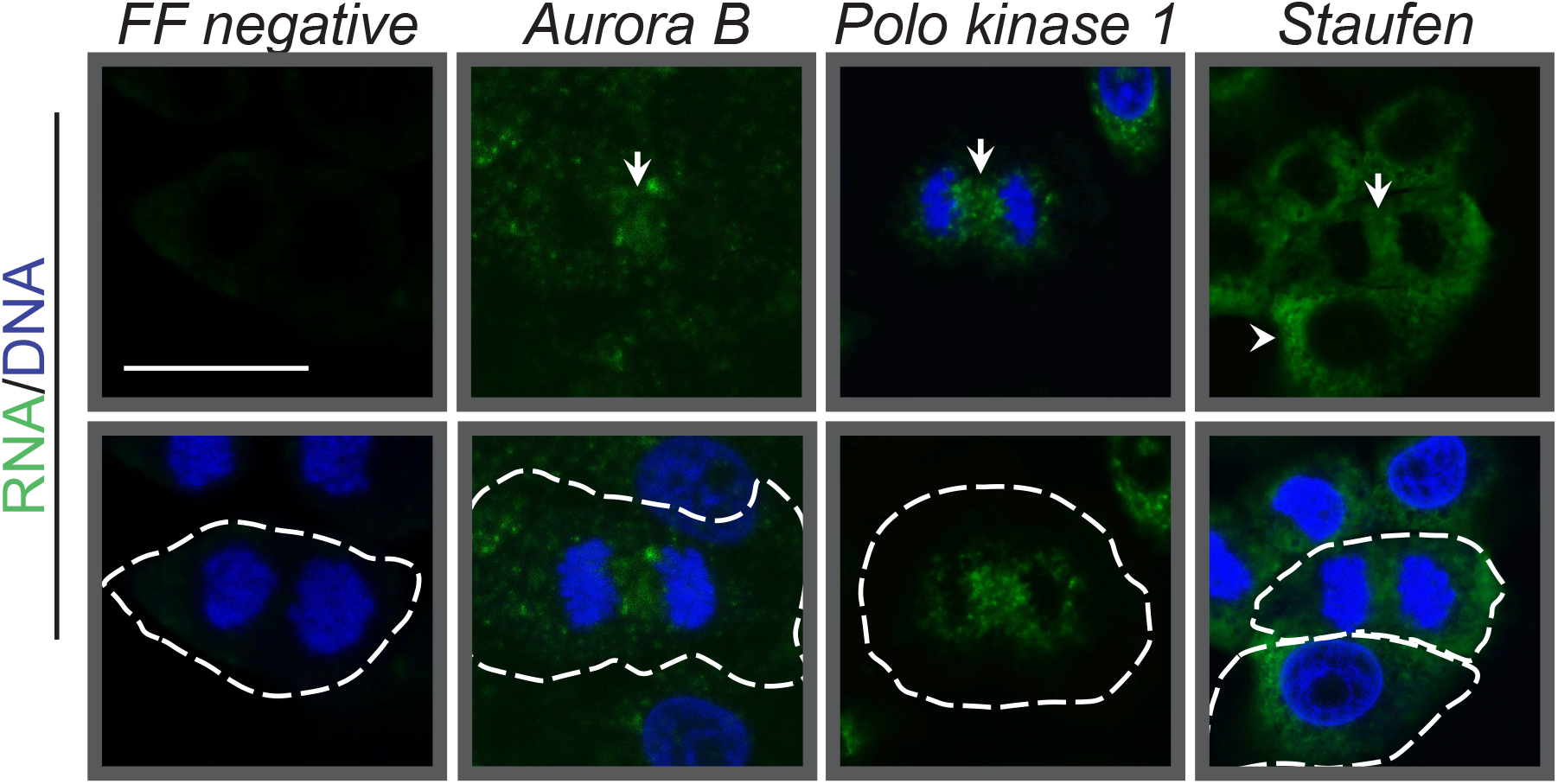
Localization of RNA transcripts is evolutionarily conserved in mammalian cells. LLC-PK1 cells were subjected to FISH, then counterstained with DAPI. Arrows indicate RNA localization at the mitotic spindle, arrowheads indicate RNA at the perinuclear region. Dashed line indicates cell boundary. FF is used as a negative control. Scale bar = 10 μm.

### The localization of RNA transcripts to the mitotic spindle is not dependent upon actin dynamics

To understand how RNA transcripts are transported to the mitotic spindle, we used cytochalasin D (Lane *et al*., 1993) to disrupt actin dynamics within the embryo, followed by examining the subcellular localization of specific transcripts. We found that the cytochalasin D resulted in an inability of the embryos to undergo cell division in a dose-dependent manner (Fig. S1A). However, cytochalasin D disruption of actin dynamics did not result in a change in localization of *Aurora B*, *Dynein, Staufen* or *Tubulin α1* transcripts (Fig. 3A). To examine this quantitatively, we measured the mean fluorescence intensity at the midzone of an anaphase blastomere and the cytoplasm and took the ratio of these measurements (Fig. 3B). A higher ratio indicates more enrichment at the mitotic spindle midzone, while a ratio of 1 indicates that the transcript is evenly dispersed throughout the blastomere. Between 2 and 8 blastomeres for each transcript were measured, and a similar trend was observed for each transcript. As a group of transcripts, we found no significant difference in the ratio of fluorescence at the spindle to the cytoplasm between embryos treated with DMSO and embryos treated with cytochalasin D (Fig. 3B). This suggests that the transport of these RNA transcripts to the mitotic spindle is not dependent upon short-term (<30 minutes) disruption of actin dynamics. In order to prevent interference with cytokinesis, we did not treat embryos with cytochalasin D for longer than 30 minutes.

**Figure 3:**
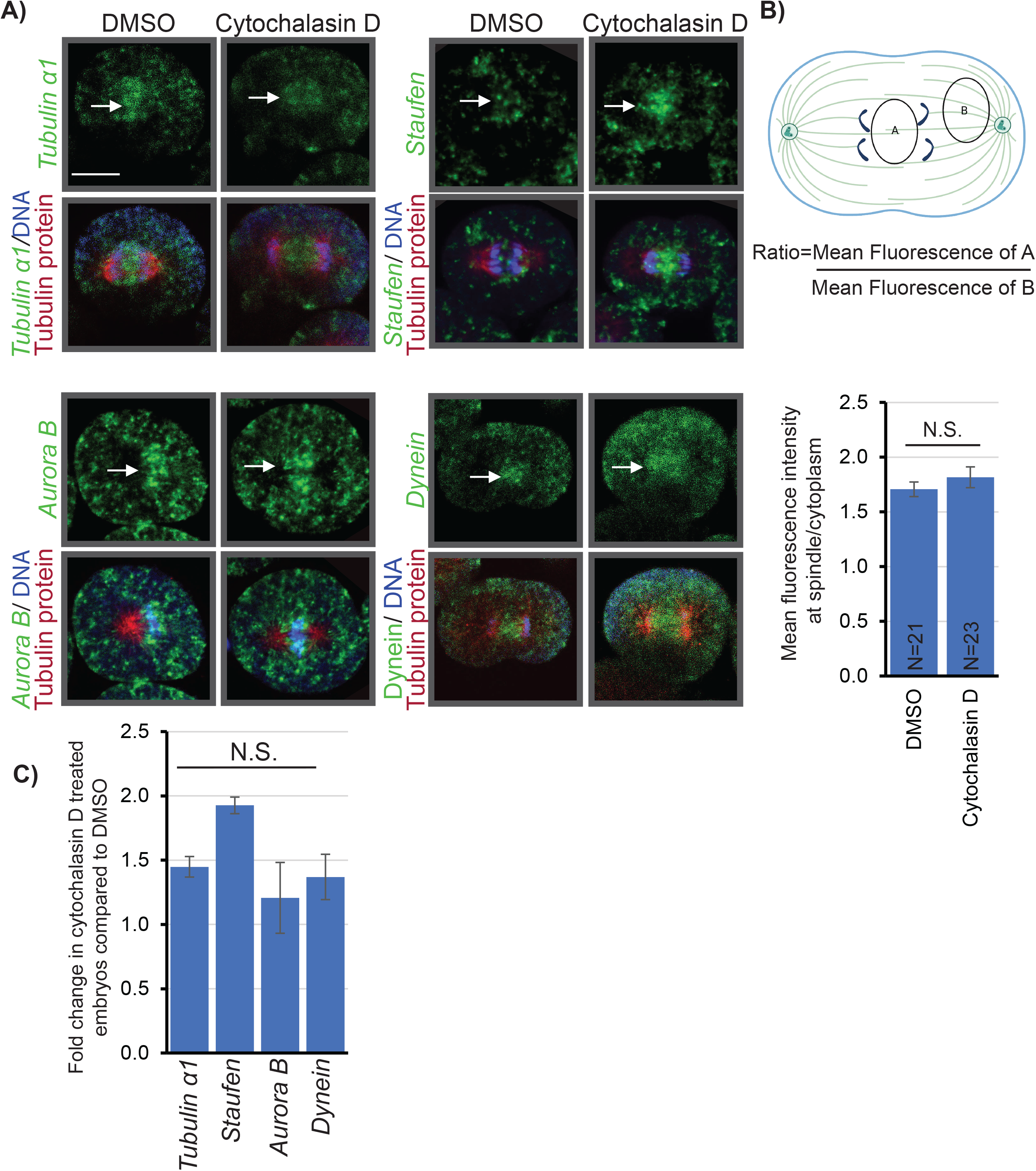
Disruption of actin polymerization with cytochalasin does not change RNA localization to the mitotic spindle. (A) RNA localization to the mitotic spindle is not affected with cytochalasin D treatment. Images are of single blastomeres of embryos at the 16-32 cell stage that were subjected to FISH with RNA probes that encode proteins known to regulate mitosis, then immunolabeled with β-tubulin antibody, and counterstained with DAPI. Arrows indicate areas of RNA localization. Scale bar = 10 μm. (B) A schematic of measured areas is shown, where area A represents a spindle region and area B represents a cytoplasmic region. The ratios of areas A and B represents the mean fluorescence intensity of RNA at the spindle to the cytoplasm. The ratio of mean fluorescence intensity of the RNA at the mitotic spindle to the cytoplasm is unchanged in embryos treated with cytochalasin D compared to control embryos. Between 2 and 8 blastomeres for each transcript were measured and a similar trend was observed for each transcript. 23 control blastomeres were measured and 21 cytochalasin D-treated blastomeres were measured. NS=no significant difference using a Student’s t-test. Error bars represent SEM. (C) qPCR results indicate no difference in transcript level between control embryos and embryos treated with cytochalasin D. 3 biological replicates.

Additionally, we observed no significant difference in transcript level between embryos treated with DMSO and embryos treated with cytochalasin D, using real time, quantitative PCR (qPCR) (Fig. 3C).

### Inhibition of microtubule polymerization disrupts localization of select RNA transcripts to the mitotic spindle

To test if localization of RNA transcripts to the mitotic spindle is dependent upon intact microtubule fibers, we used colchicine to inhibit microtubule polymerization (Rieder and Palazzo, 1992), followed by detection of the subcellular localization of specific RNA transcripts. Colchicine disrupts formation of the mitotic spindle (as observed by the immunolabeling of tubulin), as well as resulting in a more diffuse distribution of *Aurora B*, *Dynein*, *Polo kinase* and *Tubulin α1* transcripts compared to embryos treated with DMSO (Fig. 4). Treatment with colchicine results in a significantly lower ratio of fluorescence at the spindle compared to the cytoplasm in embryos treated with colchicine compared to control embryos (Fig. 4B), indicating that microtubule polymerization is required for localization of these RNA transcripts to the mitotic spindle. While the subcellular localization of these transcripts has altered, we observed no significant difference in transcript level in control embryos compared to embryos treated with colchicine, using qPCR (Fig. 4C).

**Figure 4:**
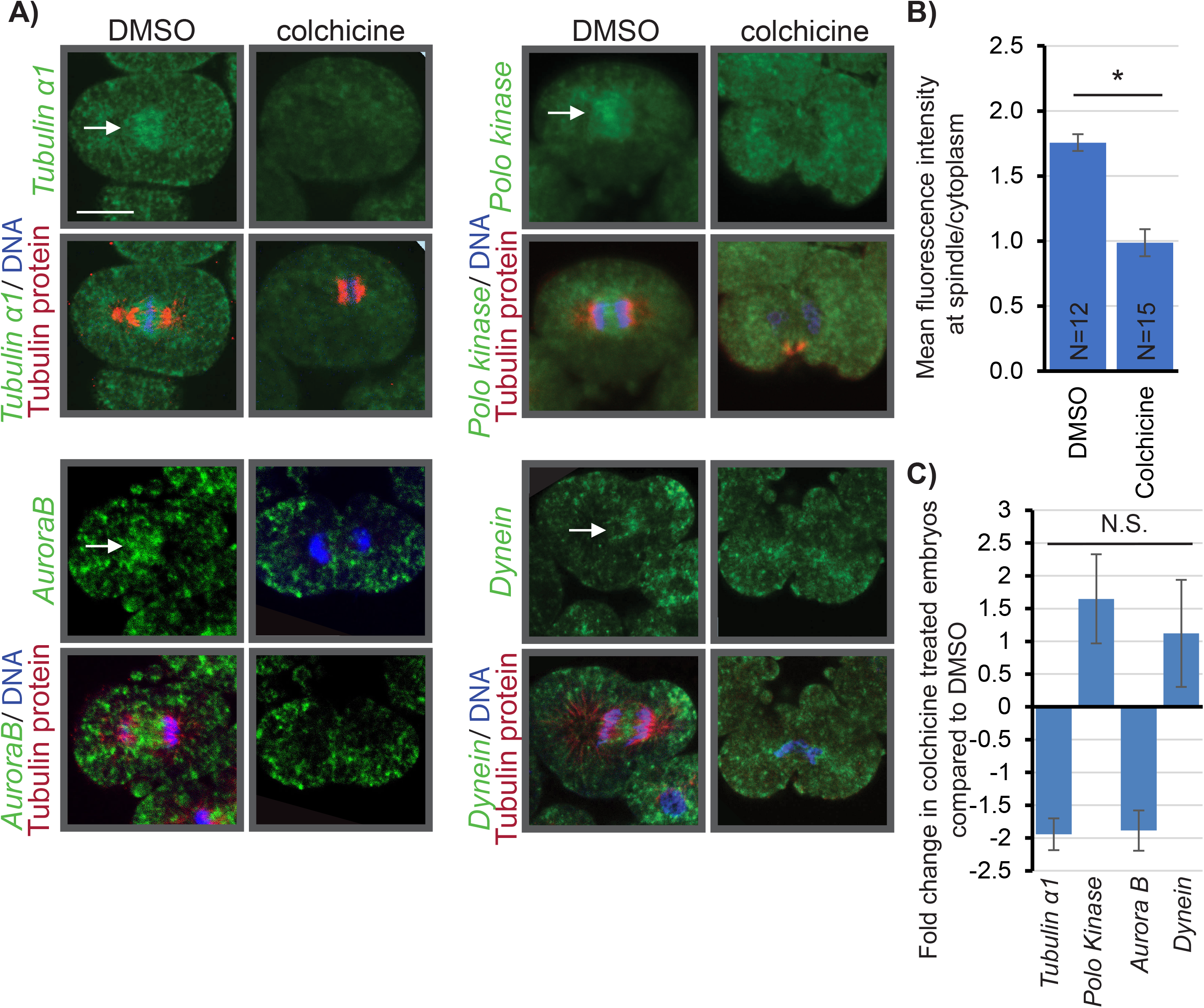
Disruption of microtubule polymerization with colchicine abrogates RNA localization to the mitotic spindle. (A) RNA is no longer localized to the mitotic spindle after embryos are treated with colchicine. Images are of single blastomeres of embryos at the 16-32 cell stage that were subjected to FISH with RNA probes of genes known to regulate mitosis, then immunolabeled with β-tubulin antibody, and counterstained with DAPI. Arrows indicate areas of RNA localization. Scale bar = 10 μm. (B) The ratio of mean fluorescence intensity of the RNA at the mitotic spindle to the cytoplasm is significantly lower in embryos treated with colchicine compared to control embryos. Between 2 and 8 blastomeres for each transcript were measured and a similar trend was observed for each transcript. 15 control blastomeres were measured and 12 colchicine blastomeres were measured. * p<0.01 using a Student’s t-test. Error bars represent SEM. (C) qPCR results indicate no difference in transcript level between control embryos and embryos treated with colchicine D. 3 biological replicates.

### Preventing kinesin-1 from interacting with its cargo results in reduced localization of RNA transcripts to the mitotic spindles

Since our data suggest that RNAs are transported to the mitotic spindle along microtubule fibers, and typically RNAs are transported subcellularly by interacting with microtubule motors (Tekotte and Davis, 2002; Suter, 2018), we investigate the role of microtubule motors in localizing RNA transcripts to the mitotic spindle. Kinesin-1 is a conserved motor protein that is known to transport vesicles, organelles and ribonucleic proteins (RNP) complexes along microtubules (Hirokawa *et al*., 2009). Kinesin-1 has been identified to regulate the localization of RNA transcripts in *Drosophila* oogenesis, neurons and cardiomyocytes (Dimitrova-Paternoga *et al*., 2021; Fukuda *et al*., 2021; Scarborough *et al*., 2021). We used kinesore, a drug which binds to kinesin-1 at the cargo site and activates kinesin 1’s ability to bind to microtubules (Randall *et al*., 2017). RNA localization to the mitotic spindles in embryos treated with kinesore is significantly reduced compared to embryos treated with DMSO (Fig. 5A,B), as indicated by a statistically significant reduction of the ratio of mean fluorescence intensity at the mitotic spindle to the mean fluorescence intensity in the cytoplasm (Fig. 5B). This suggests that these RNAs are in part transported to the mitotic spindle by kinesin-1. We found that the total level of these transcripts was not significantly different in embryos with kinesore compared to embryos treated with DMSO (Fig. 5C).

**Figure 5:**
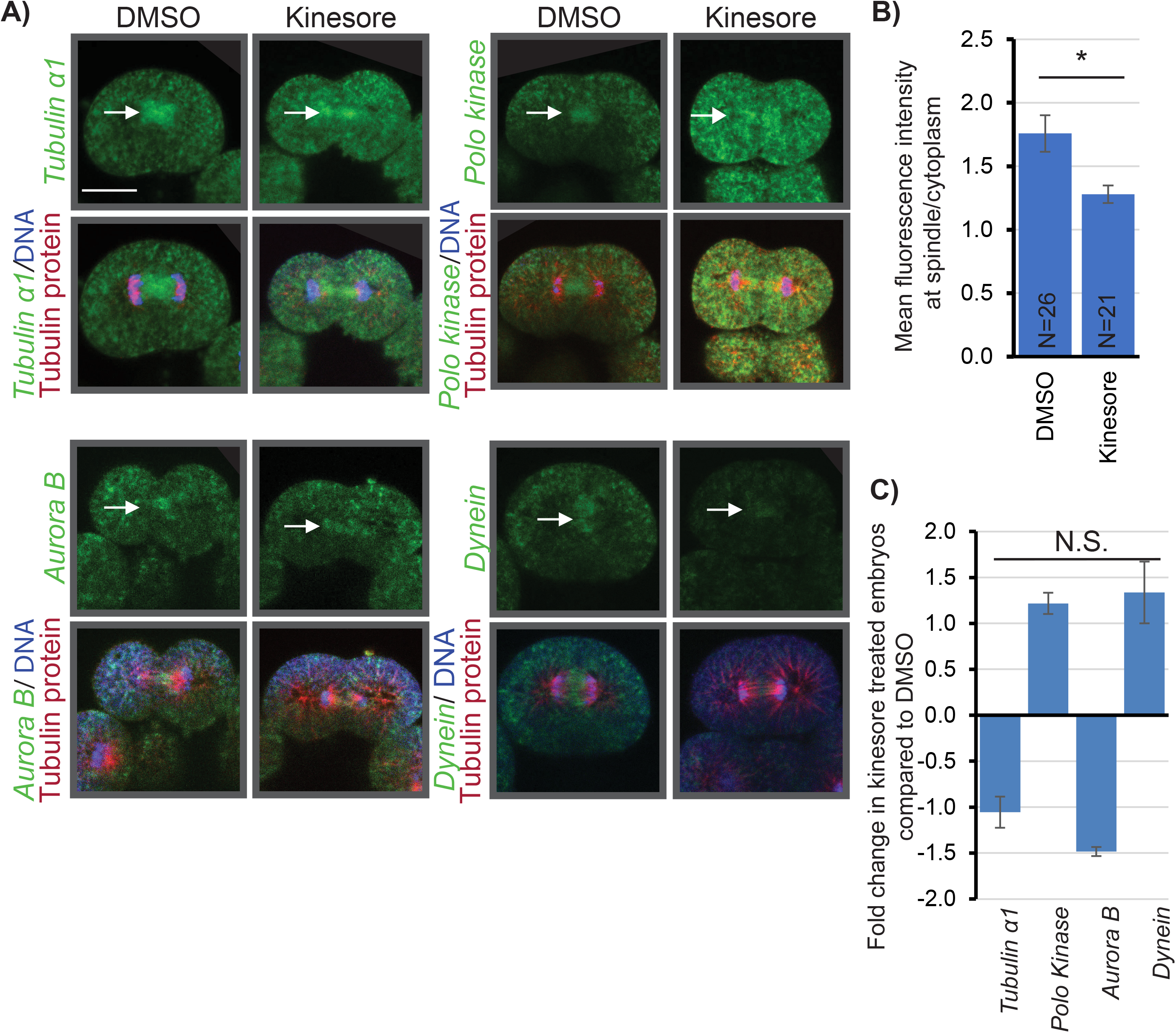
Kinesore treatment diminishes RNA localization to the mitotic spindle. (A) RNA is less localized to the mitotic spindle after embryos are treated with kinesore. Images are of single blastomeres of embryos at the 16-32 cell stage that were subjected to FISH with RNA probes of genes known to regulate mitosis, immunolabeled with β-tubulin antibody, and counterstained with DAPI. Arrows indicate areas of RNA localization. Scale bar = 10 μm. (B) The ratio of mean fluorescence intensity of the RNA at the mitotic spindle to the cytoplasm is significantly lower in embryos treated with kinesore compared to control embryos. Between 2 and 8 blastomeres for each transcript were measured and a similar trend was observed for each transcript. 26 control blastomeres were measured and 21 kinesore blastomeres were measured. * p<0.01 using a Student’s t-test. Error bars represent SEM. (C) qPCR results indicate no difference in transcript level between control embryos and embryos treated with kinesore. 3 biological replicates.

### Preventing dynein from transporting its cargo along microtubules alters localization of RNA transcripts at the mitotic spindle

As kinesin-1 is a plus-ended motor (Hirokawa *et al*., 2009), we also wanted to examine dynein, a minus-ended motor that has been identified to transport RNA transcripts in *Drosophila* oogenesis and neurons (Schnorrer, Bohmann and Nusslein-Volhard, 2000; Xu, Brechbiel and Gavis, 2013; Herbert *et al*., 2017). We used ciliobrevin D to inhibit dynein from transporting its cargo along the microtubule filaments (Firestone *et al*., 2012). As has been reported previously, we observe smaller and more compact mitotic spindles in ciliobrevin-treated embryos compared to DMSO treated embryos (Fig. 6A, tubulin immunolabeling) (Firestone *et al*., 2012).

**Figure 6:**
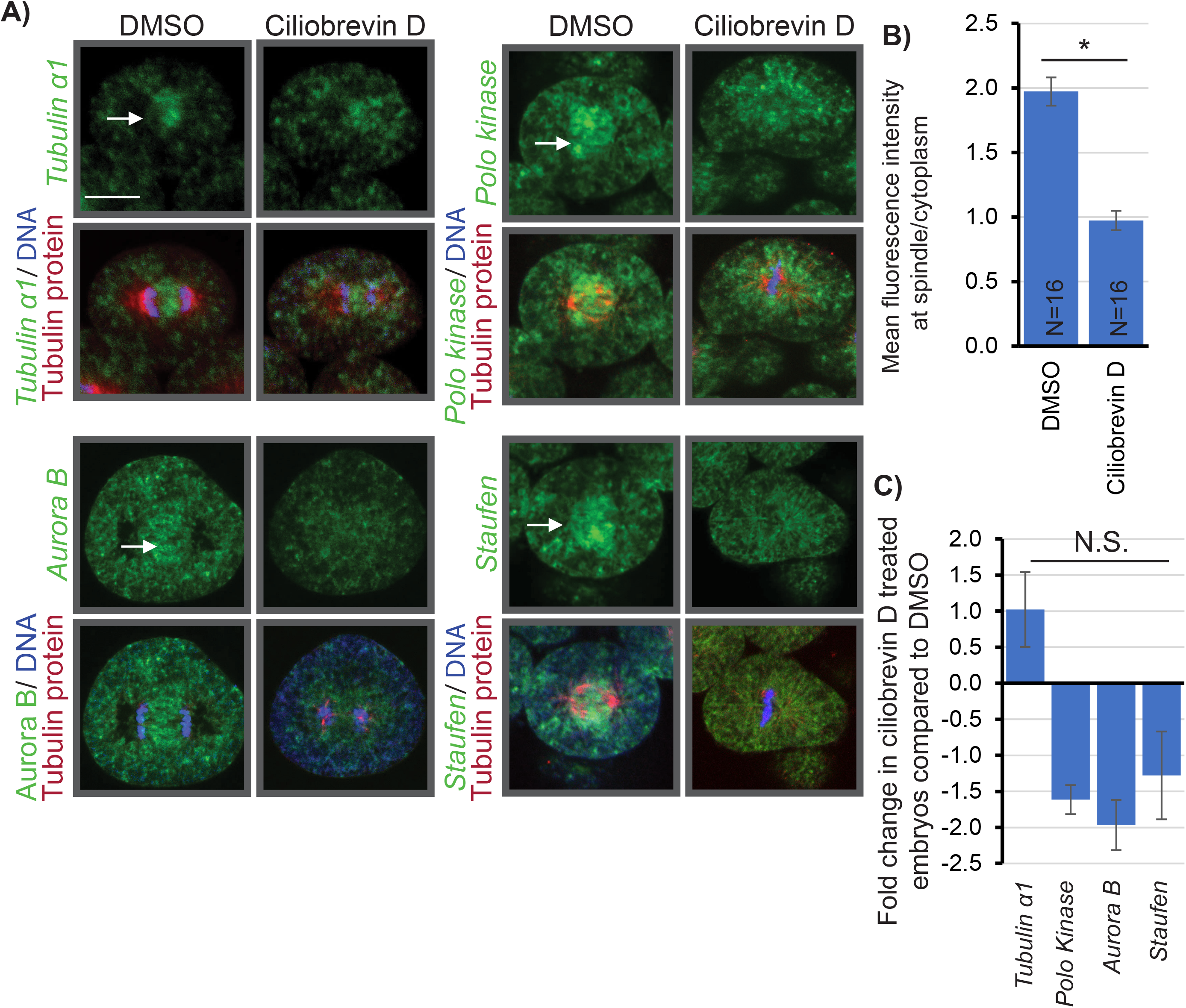
Dynein inhibition alters RNA localization to the mitotic spindle. (A) RNA localization to the mitotic spindle is altered after embryos are treated with ciliobrevin D. Images are of single blastomeres of embryos at the 16-32 cell stage that were subjected to FISH with RNA probes of genes known to regulate mitosis, immunolabeled with β-tubulin antibody, then counterstained with DAPI. Arrows indicate areas of RNA localization. Scale bar = 10 μm. (B) The ratio of mean fluorescence intensity of the RNA at the mitotic spindle to the cytoplasm is significantly lower in embryos treated with ciliobrevin D compared to control embryos. Between 2 and 8 blastomeres for each transcript were measured and a similar trend was observed for each transcript. 16 control blastomeres were measured, 16 ciliobrevin D blastomeres were measured. * p<0.01 using a Student’s t-test. Error bars represent SEM. (C) qPCR results indicate no difference in transcript level between control embryos and embryos treated with ciliobrevin D. 3 biological replicates.

In control embryos, RNA transcripts are enriched at the midzone of the mitotic spindle, while in ciliobrevin D treated embryos, the RNA transcripts are enriched at the plus ends of the astral microtubule filaments (Fig. 6A). The ratio of fluorescence at the spindle compared to the cytoplasm is significantly decreased in embryos treated with ciliobrevin D compared to embryos treated with DMSO, indicating that transport of the RNA transcripts to the spindle is dependent on dynein (Fig. 6B). Despite changes of *Staufen*’s subcellular localization, we observe no change in the level of *Staufen* in embryos treated with ciliobrevin D compared to control embryos, using qPCR (Fig. 6C).

### The CPE within the 3’UTR of *Aurora B* is necessary for the localization of *Aurora B* RNA transcript to the mitotic spindle and critical for early development

In order to understand how RNA transcripts are localized to the mitotic spindle, we investigated which region of the transcript is necessary for localization. We cloned the 3’UTR of *Aurora B* downstream of *Renilla* luciferase (*Rluc*) construct (*Aurora B-Rluc*) and tested its localization within the dividing embryo (Fig. 7A,B). Results indicate that the 3’UTR is necessary and sufficient to localize *AuroraB-RLuc* to the mitotic spindles, while *Rluc* transcript by itself does not localize to the mitotic spindles (Fig. 7A,B). We bioinformatically identified a potential CPE within its 3’UTR. To test if the CPE within *Aurora B* is critical for its localization to the mitotic spindles, we deleted the CPE in the *Aurora B* 3’UTR downstream of *RLuc* (Fig. 7A,B). Results indicate that deletion of the CPE abrogates localization of the *AuroraB-RLuc* transcript at the mitotic spindles (Fig. 7A).

**Figure 7:**
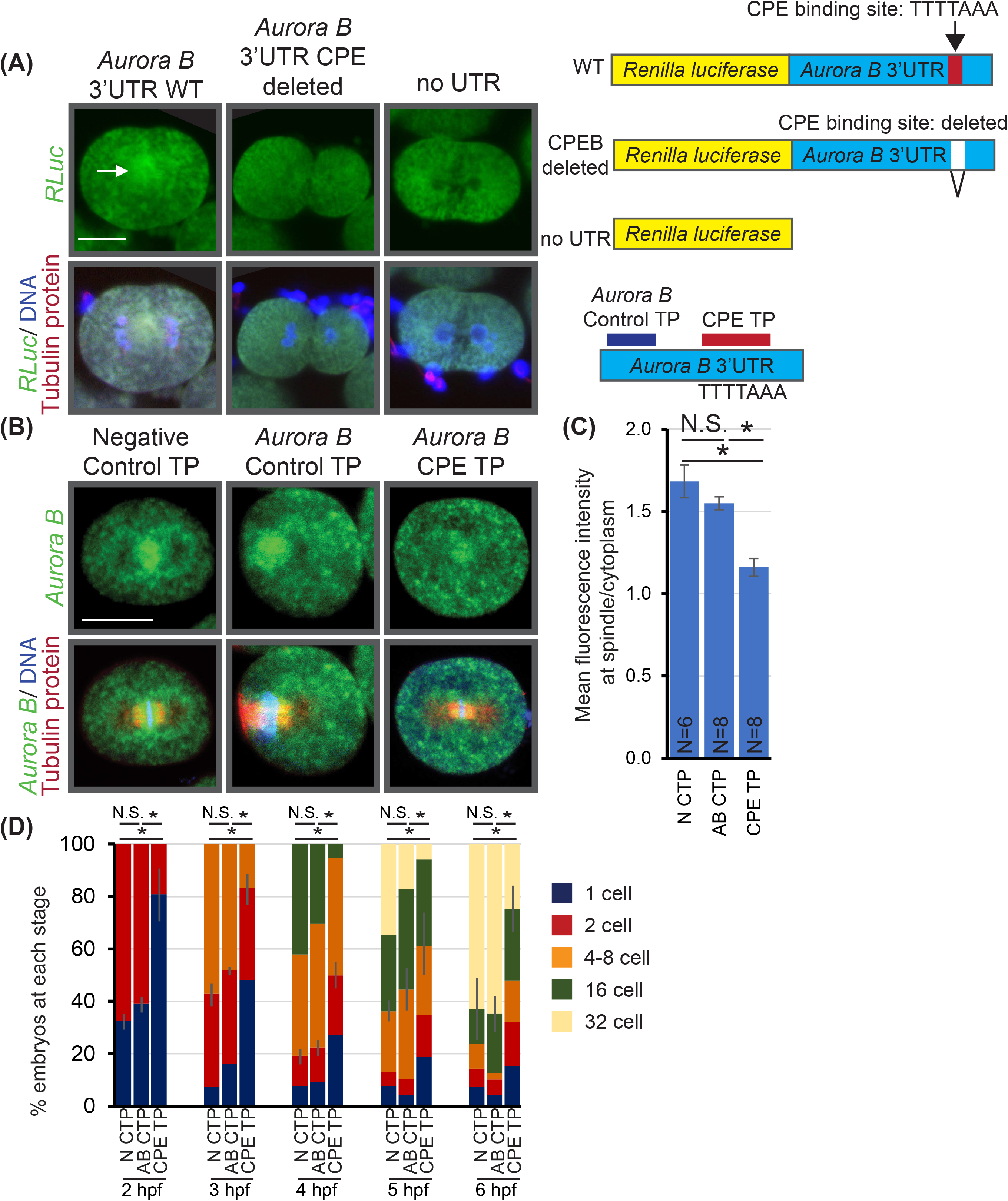
The 3’UTR of *Aurora B*, and specifically the CPE sequence, is necessary for localization of RNA to the mitotic spindle. (A) Embryos were injected with RNA constructs containing *RLuc-Aurora B 3’UTR WT* RNA, *RLuc-Aurora B 3’UTR CPE deleted*, or *RLuc-no UTR.* Exogenously injected *RLuc-Aurora B 3’UTR WT* RNA is localized at the mitotic spindle, while *RLuc-Aurora B 3’UTR CPE* deleted is no longer localized, similar to *RLuc-no UTR* RNA. Images are of single blastomeres of embryos collected at the 16-32 cell stage then subjected to FISH with the *RLuc* probe, immunolabeled with β-tubulin antibody, then counterstained with DAPI. Arrows indicate areas of RNA localization. Scale bar = 10μm. (B) Blocking the CPE within the 3’UTR of *Aurora B* results in less localization of endogenous *Aurora B* to the mitotic spindle compared to control embryos. Images are of single blastomeres of embryos collected at the 16-32 cell stage then subjected to FISH *Aurora B* RNA probe, immunolabeled with β-tubulin antibody, then counterstained with DAPI. Arrows indicate areas of RNA localization. Scale bar = 10 μm. (C) The ratio of mean fluorescence intensity of the RNA at the mitotic spindle to the cytoplasm is significantly lower in embryos in which the CPE is blocked compared to control embryos. 6 negative control blastomeres, 8 *Aurora B* 3’UTR control TP blastomeres and 8 *Aurora B* CPE TP blastomeres were measured. * p<0.005 using a one-way ANOVA, with a post-hoc Tukey-Kramer test . Error bars represent SEM. (D) Embryos in which the CPE is blocked have significant developmental delay compared to control embryos. *p<0.001 using Cochran-Mantel-Haenszel test. 210 negative control embryos, 255 *Aurora B 3’UTR* control embryos, 217 *Aurora B* CPE TP embryos, 3 biological replicates. Error bars indicate SEM.

To test if localization of *Aurora B* RNA to the mitotic spindle has an impact on embryonic development, we designed a synthetic morpholino antisense oligonucleotide complementary to the CPE to block potential binding of CPEB to the endogenous CPE within the *Aurora B* 3’UTR (Fig. 7B). Results indicate that blocking the CPE significantly reduces localization of endogenous *Aurora B* RNA to the mitotic spindles (Fig. 7C), as the ratio of fluorescence at the spindle compared to the cytoplasm is significantly decreased in embryos in which the CPE is blocked compared to control embryos (Fig. 7C).

Importantly, blocking the *Aurora B* CPE results in early developmental defects. This is evident that as early as 2 hpf, 61% of embryos injected with the control oligo have divided to 2 cells, whereas only 19% of embryos injected with CPE blocking oligo have divided to 2 cells (Fig. 7D). This trend persists throughout the early cleavage stages to 6 hpf, where 64.7% of embryos injected with the control oligo have reached the 16-32 cell stage, compared to 24.8% of the embryos injected with CPE blocking oligo have reached the same developmental stage (Fig. 7D). There is no significant difference between embryos injected with the negative control oligo, which does not recognize specific sequences within the sea urchin genome, and the *Aurora B* control oligo, which is complementary to the *Aurora B* 3’UTR sequence upstream of the CPE. Of note is that at 24 hpf, only 50% of the embryos injected with CPE blocking oligo have developed into blastulae, compared to 85% of injected control embryos (Fig S2). Taken together, since 50% of CPE blocking oligo-injected embryos do not survive to 24 hpf, mislocalization of *Aurora B* transcript likely lead to developmental arrest or lethality. Thus, these data indicate that localization of *Aurora B* transcript to the mitotic spindle is important for early development.

## Discussion

During early development, cells go through rapid cycles of mitosis, without intervening gap phases, making regulation of mitosis critical during this time (Siefert, Clowdus and Sansam, 2015). We identified RNA localization as a potential regulatory mechanism that regulates the relatively fast cell divisions during the cleavage stage embryos. Biochemical assays have identified transcripts that regulate cell cycle, cell division and chromosome function to be enriched in the subset of transcripts associated with mitotic spindles (Sharp *et al*., 2011). These assays were performed in *Xenopus* egg extract in which mitotic spindle formation was induced. To date, only a select few transcripts, such as *cyclin B* and *vasa*, have been visualized at the mitotic spindle in developing embryos (Groisman *et al*., 2000; Sharp *et al*., 2011; Yajima and Wessel, 2015; Takahashi, Ishii and Yamashita, 2018; Waldron and Yajima, 2020; Fernandez-Nicolas *et al*., 2022).

We observed that transcripts encoding proteins involved in mitosis localize to the spindles (Fig. 1A,B), while transcripts encoding proteins that do not regulate mitosis do not show that localization (Fig. 1C). Prior research has indicated that localization of the RNA correlates with the site where the encoded protein functions (Mowry and Melton, 1992; Kloc and Etkin, 1994; Höfer, Ness and Drenckhahn, 1997; Joseph and Melton, 1998; Farina *et al*., 2003). For example, *Vg1* mRNA and protein localizing to the vegetal pole of the *Xenopus* oocyte, where the localization of *Vg1* mRNA is known to be important for inducing endoderm and mesoderm in developing *Xenopus* embryos (Mowry and Melton, 1992; Kloc and Etkin, 1994; Joseph and Melton, 1998). Another example is that disruption of *β-actin* mRNA and protein localization to lamellipodia in chicken fibroblasts alters the polarization and migration of the cell (Höfer, Ness and Drenckhahn, 1997; Shestakova, Singer and Condeelis, 2001; Farina *et al*., 2003). In addition, intracellular RNA localization has been well studied in the context of neurons (Mayford *et al*., 1996; Huang *et al*., 2003; Dahm, Kiebler and Macchi, 2007; Yoon *et al*., 2016, and many others). Neurons can have extremely long axons, and contain distinct intracellular regions, such as dendrites and synaptic boutons that have markedly different local environments (Li, Yu and Ji, 2021). The different local environments within different parts of a neuron are partially due to local translation of transcripts, such as *CamKIIα* and *MAP2*, which is thought to regulate synaptic activity and neuronal plasticity (Huang *et al*., 2003; Dahm, Kiebler and Macchi, 2007; Doyle and Kiebler, 2011). These examples highlight the functional importance between the localization of transcripts and the ultimate localization of their corresponding proteins.

Subcellular RNA localization has been identified to occur through a molecular motor transporting the RNA along a cytoskeletal element, such as actin and microtubule filaments (Tekotte and Davis, 2002). For example, RNAs known to be dependent upon actin for localization include *Ash1* in budding yeast (Takizawa *et al*., 1997; Beach and Bloom, 2001), and *β-actin* in embryonic fibroblasts (Latham *et al*., 2001), and *MAP2* in neurons (Balasanyan and Arnold, 2014). However, we found that while short-term disruption of actin dynamics by cytochalasin D disrupts development (Fig. S1A), it does not alter localization of RNA transcripts at the mitotic spindles (Fig. 3A, B). This result suggests that transport of majority of these RNAs is not along actin filaments, or that the level of actin disruption was insufficient to disrupt RNA localization.

Of note is that in order to quantify changes in localization of RNA transcripts at the spindle, we utilized a ratio of the mean fluorescence of a region between the dividing nuclei at anaphase to the mean fluorescence of an identically sized region in the cytoplasm. Importantly, the ratio of fluorescence at the spindle compared to the cytoplasm is similar among the DMSO-treated controls for all the small molecule inhibitors used (Fig. 3B, 4B, 5B and 6B), as well as between the control injected embryos (Fig. 7C). This indicates that this is a consistent way to objectively measure RNA localization to the spindle.

We observed that disruption of microtubule dynamics by colchicine abrogated the localization of these RNA transcripts at the mitotic spindle (Fig. 4A, B). This localization of transcripts could be due to RNA transcripts being actively transported along the microtubule filaments in a complex with a motor protein, as has been observed in neurons, muscles and fly embryos, among others (Lyons *et al*., 2009; Goldman and Gonsalvez, 2017; Denes, Kelley and Wang, 2021). For example, the RNA-binding protein CPEB, known to localize *MAP2* to dendrites, has been found in granules with the motor proteins dynein and kinesin, suggesting that transport may occur along microtubule tracks (Huang *et al*., 2003). Alternatively, the disruption of localization of these RNA transcripts upon colchicine treatment could be due to the RNA transcripts being anchored at the spindle and disruption of the microtubule filaments results in passive diffusion of the RNA transcripts. For example, apically localized transcripts in *Drosophila* blastoderm embryos, such as *run* and *ftz* transcripts are transported to the apical end of the embryos by dynein, which then becomes anchored to sponge bodies, which are electron-dense particles related to P-bodies, in a microtubule dependent manner (Delanoue and Davis, 2005). Disruption of microtubule dynamics alters localization of RNA to the mitotic spindle (Fig. 4A, B), indicating RNA localization to the spindle is dependent upon intact microtubules, but does not distinguish between transport of RNA along the microtubules and anchoring of RNA to the microtubules.

To identify whether RNAs are transported along microtubule filaments, we investigated the role of motor proteins. The main motors that have been implicated in RNA transport are myosin, which transports RNA along actin filaments, and kinesins and dynein, which transport RNA along microtubule filaments (Takizawa *et al*., 1997; Januschke *et al*., 2002; Tekotte and Davis, 2002; Messitt *et al*., 2008; Xu, Brechbiel and Gavis, 2013). To identify the role of microtubule motors in localization of RNA transcripts to the mitotic spindle, we used kinesore, a small-molecule inhibitor which prevents kinesin-1 from binding to its cargo (Randall *et al*., 2017). We observed reduced localization of transcripts to the mitotic spindle in kinesore-treated embryos compared to control embryos (Fig. 5A, B). Our result is consistent with prior literature in which kinesin-1 has been identified to localize several RNA transcripts, such as *oskar* in *Drosophila* oocytes, *CamKIIα* in oligodendrocytes (Kanai, Dohmae and Hirokawa, 2004), and *cyclin B* in *Danio* oocytes (Takahashi, Ishii and Yamashita, 2018). Both kinesin-1 and kinesin-2 are required to localize *Vg1* in *Xenopus* oocytes (Messitt *et al*., 2008). Our data indicate that kinesin-1 plays a role in the localization of RNA transcripts to the mitotic spindles. We observe that kinesore has very little impact on the overall levels of transcripts, based on our qPCR data (Fig. 5C). While RNA *in situ* using FISH suggests that there may be slightly less *Dynein* in kinesore-treated embryos compared to control, the qPCR shows a small decrease, with no statistical significance. One caveat is that while qPCR analysis is quantitative, the embryos collected for this analysis are not all undergoing mitosis. Thus, if *Dynein* or another transcript undergoes cell-cycle specific changes in expression, this would not be detected with qPCR. However, the focus here is to examine the spatial localization of transcripts and we found kinesin-1 to be important for RNA localization.

Inhibition of AAA ATPase of dynein with ciliobrevin D results in transcript accumulation to the plus ends of the astral microtubules (Fig. 6A). This may be due to dynein directly transporting the RNA or through the ability of dynein to anchor RNA to microtubules (Delanoue and Davis, 2005). In addition, we observed that the mitotic spindle appears smaller (Fig. 6A), which has been observed previously in ciliobrevin D-treated cells (Firestone *et al*., 2012). The exact mechanism of the smaller spindle is not known, but this may be due to the dynein’s role in anchoring astral microtubules to the cortex of the cell (Hueschen *et al*., 2017), and its ability to mediate microtubule sliding in a cortical direction (Okumura *et al*., 2018). It is also possible that kinesin-1 and dynein cooperate to ensure proper RNA localization, as observed in *Drosophila* embryos, where both kinesin-1 and dynein work together to properly localize *bcd* and *gurken* RNAs (Januschke *et al*., 2002). Since motor proteins kinesin-1 and dynein, as well as intact microtubules are needed for the localization of RNA at the spindles, our overall results indicate that the transcripts are transported along microtubules to their final destination at the midzone of the mitotic spindle.

We identified that the 3’UTR of *Aurora B* is necessary and sufficient for its localization to the mitotic spindles (Fig. 7B). In addition, deletion of the CPE or blockage of the CPE within the 3’UTR prevents localization of exogenous *Aurora B-Rluc* to the mitotic spindles (Fig. 7B,C). CPEB has been identified to be necessary for localization of *cyclin B* RNA of *Xenopus* embryos (Groisman *et al*., 2000), as well as for *BUB3* RNA, which encodes a mitotic checkpoint protein, to the mitotic spindle (Pascual *et al*., 2021). Preventing CPEB from binding to *cyclin B* RNA results in defects in mitosis (Groisman *et al*., 2000; Pascual *et al*., 2021). Similar to *cyclin B*, deletion of CPE within *Aurora B* 3’UTR had completely abolished *Aurora B*’s localization (Fig. 7A). Further, blocking CPE site within *Aurora B* 3’UTR also resulted in a significant reduction of localized transcript at the spindles (Fig. 7B-C). This reduction, rather than a complete abolishment of localization, may be due to the AT-rich sequence in the 3’UTR region that result in a weaker binding of the blocking oligo to the endogenous CPE within *Aurora B* transcripts. Importantly, we also observed that blocking the CPE in endogenous *Aurora B* transcript results in developmental arrest (Fig. 7D). Approximately 50% of the CPE blocking oligo-injected embryos do not live to the blastulae stage (24 hpf) (Fig S2), indicating that the embryos experiencing developmental arrest do not survive. We do not know the exact mechanism of how mislocalization of *Aurora B* transcript away from the spindles causes developmental arrest and lethality. Potentially, disrupting localization of *Aurora B* transcript has a similar effect as blocking *cyclin B* transcript’s localization to the mitotic spindles (Groisman *et al*., 2000). In the case of mislocalization of *cyclin B*, these mitotic defects occur while cyclin B protein levels continue to display normal oscillations throughout the cell cycle, similar to the control (Groisman *et al*., 2000). It was suggested that the defects in mitosis are not due to a global deficit of cyclin B protein, but rather, the localization of the *cyclin B* transcript and its local translation at the mitotic spindle itself is important for progression through mitosis (Groisman *et al*., 2000). Since ribosomal proteins and RNAs are present at the mitotic spindle and in early cleavage stage embryos (Hassine *et al*., 2020; Fernandez-Nicolas *et al*., 2022), an intriguing possibility is that local translation of these transcripts encoding proteins that regulate mitosis may be essential for mitotic progression.

Aurora B functions by sensing bi-polar attachment of chromosomes to the mitotic spindle (Krenn and Musacchio, 2015). This is essential to Aurora B’s regulation of the SAC through phosphorylation of its substrates leading to degradation of securin (Lens *et al*., 2003; Krenn and Musacchio, 2015). Interestingly, preventing Aurora B protein from localizing to the centrosomal region results in defects in mitosis, despite the fact that they retain kinase ability (Scrittori *et al*., 2005). Together with results from prior studies and our study, we propose that Aurora B’s protein function is tightly tied to its transcript localization which adds another layer of regulation of mitosis (Fig. 8). We propose that this regulation extends to other important mitotic regulators as well. Intriguingly, this RNA localization is a conserved phenomenon observed in mammalian cells as well (Fig. 2). Since mitosis, especially during the early cleavage stages of development, must occur rapidly and be tightly controlled, localizing the RNA of key players of mitosis may be an evolutionarily conserved mechanism to facilitate rapid changes in the translation of these RNAs, allowing for proper cell division to occur.

**Figure 8:**
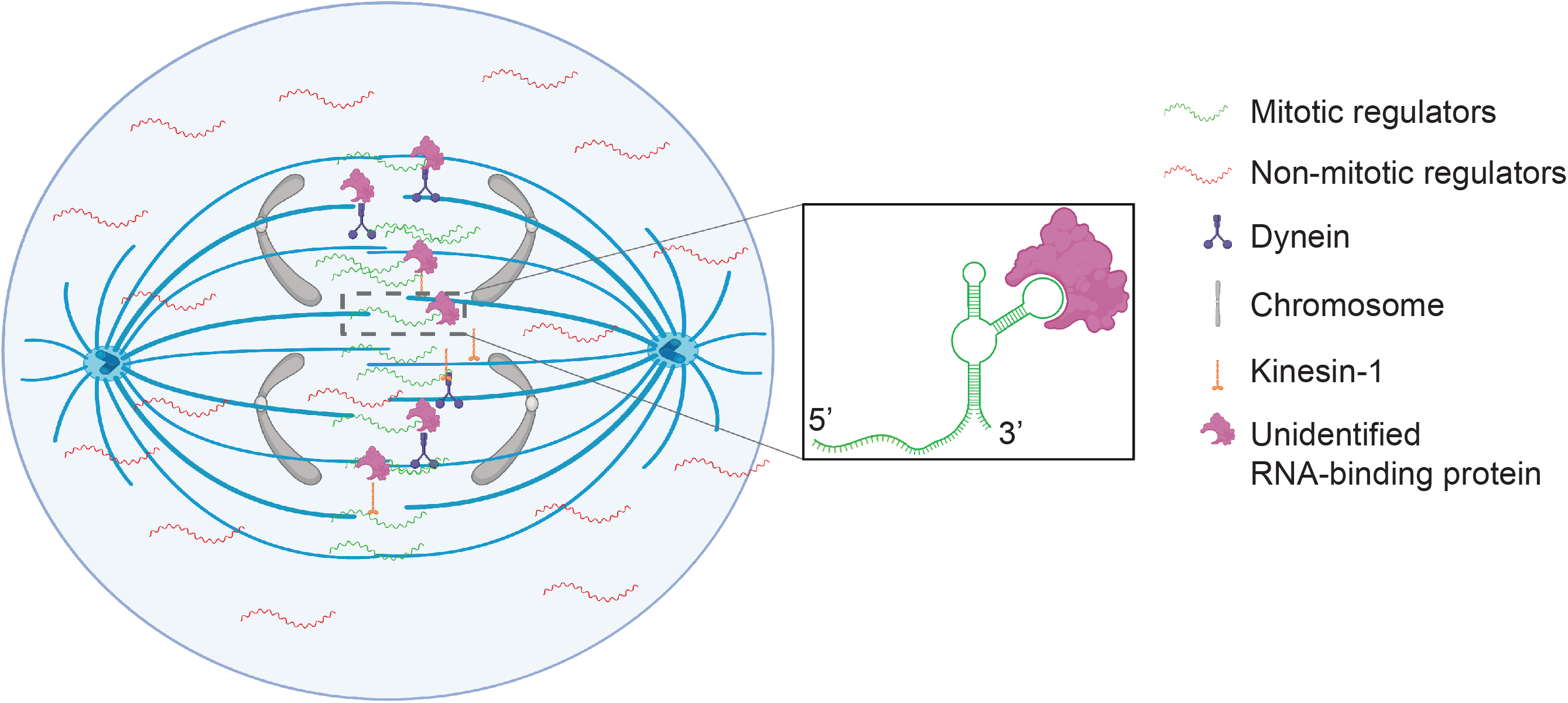
Model of RNA transcript localization to the mitotic spindle. RNA transcripts that encode proteins that regulate mitosis are localized to the mitotic spindle. Microtubule motors kinesin-1 and dynein are involved in the transport of these RNA transcripts. Based on our results as well as previous research, we hypothesize that local translation of these transcripts (*cyclin B*, *Aurora B*) are essential for proper development (Groisman *et al*., 2000). The 3’UTR of the RNAs contain sequences responsible for binding of RNA-binding proteins, such as CPEB, which may be responsible for localization of the RNA to the mitotic spindle. RNA localization allows local translation at the spindles is a regulatory mechanism to ensure rapid cell divisions that occur during the early cleavage stage.

## Materials & Methods

### Animals

Adult *Strongylocentrotus purpuratus* were collected from Marinus Scientific, LLC (Lakewood, CA) or Point Loma Marine Invertebrate Labs (Lakeside, CA) and were maintained at 12°C in artificial sea water (ASW) made from distilled, deionized water and Instant Ocean^©^. Adults were induced to shed either through shaking or intracoelomic injection of 0.5 M KCl. Embryos were cultured at 12°C in filtered natural sea water (FSW) obtained from the Indian River Inlet (University of Delaware).

### Cell culture

LLC-PK1(Hull, Cherry and Weaver, 1976) (LLC-PK1, ATCC No. CL-101) cells were maintained in DMEM/F12 (ThermoFisher Scientific, Waltham, MA) media supplemented with 10% fetal bovine serum (MilliporeSigma, Burlington, MA) at 37°C under 5% CO_2_.

### Fluorescence RNA in situ Hybridization (FISH) and immunolabeling

The steps performed for FISH are described previously with modifications (Sethi, Angerer and Angerer, 2014). RNA *in situ* hybridization probes were amplified using sea urchin cDNA for sea urchin specific probes and porcine cDNA for mammalian probes. Primers were synthesized based on known sequences (IDTDNA, Coralville, Iowa) and amplicons were ligated into the ZeroBlunt vector (ThermoFisher Scientific, Waltham, MA) (Table 1). Positive clones were sequenced (Genewiz Services, South Plainfield, NJ), digested (ThermoFisher, Scientific, Waltham, MA) and DIG labeled using specific RNA polymerases (MilliporeSigma,Burlington, MA) decribed in Table 1. Probe was used at 0.5 ng probe/µL to detect native transcript in embryos, according to previous protocols (Stepicheva *et al*., 2015). The embryos were incubated with anti-digoxigenin-POD antibody at 1:1,000 (MilliporeSigma, St.Louis, MO) overnight at 4°C and amplified with Tyramide Amplification working solution (1:150 dilution of TSA stock with 1x Plus Amplification Diluent-fluorescence) (Akoya Biociences, Marlborough, MA). The embryos were washed with MOPS buffer three times then with PBST (1xPBS,0.1% TritonX-100) three times. After FISH, embryos were incubated for overnight at 4°C in E7 antibody against β-tubulin (Developmental Studies Hybridoma Bank, Iowa City, IA) diluted to 5 μg mL^-1^ in 4% sheep serum in PBST. Embryos were washed three times with PBST then incubated for 1 hour at room temperature with secondary antibody (Alexa-Fluor 647, Thermo Fisher Scientific, Waltham, MA) diluted at 1:300 in 4% sheep serum (MilliporeSigma, Burlington, MA) in PBST. Embryos were washed three times with PBST, then counter-stained with DAPI (ThermoFisher Scientific, Waltham, MA). Images were obtained with a Zeiss LSM 780 or 880 scanning confocal microscope (Carl Zeiss Incorporation, Thorwood, NY). Single digital image or the maximum intensity projections of Z-stack of images were acquired with Zen software and exported into Adobe Photoshop and Illustrator (Adobe, San Jose, CA) for further processing.

**Table 1:**
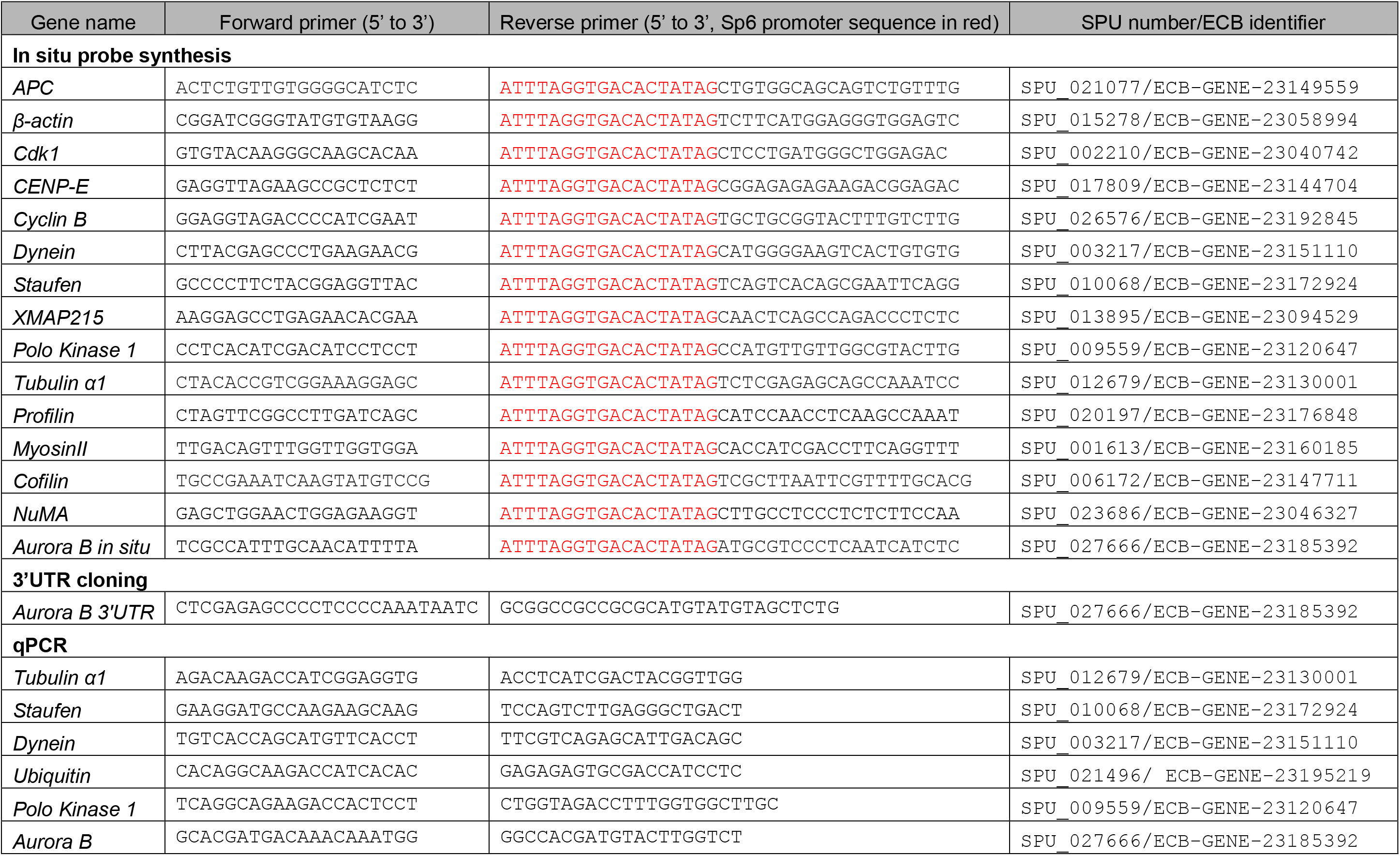
Primers used in this study

LLC-PK cells were fixed (100 µM MOPS, 0.1% Tween, 4% paraformaldehyde in PBS) for 15 minutes at room temperature, then washed with PBS 0.01%Tween. Cells were hybridized as described (Martín-Durán *et al*., 2017), and incubated with 0.5 ng µL^-1^ probe at 50°C for 48 hours. The cells were incubated with anti-digoxigenin-POD antibody at 1:1,000 (MilliporeSigma, Burlington, MA) for 1 hour at room temperature and amplified with Tyramide Amplification working solution (1:150 dilution of TSA stock with 1x Plus Amplification Diluent-fluorescence) (Akoya Biosciences, Marlborough, MA). Cells were mounted in VectaShield Anti-Fade mounting media with DAPI (Vector Laboratories, Newark, CA). Images were obtained with a Zeiss LSM 780 or 880 scanning confocal microscope (Carl Zeiss Incorporation, Thorwood, NY). Single digital image or the maximum intensity projections of Z-stack of images were acquired with Zen software and exported into Adobe Photoshop and Illustrator (Adobe, San Jose, CA) for further processing. Excess DNA in the images is due to sperm and does not affect the interpretation of the results.

### Microinjections and RNA constructs

Microinjections were performed as previously described with modifications (Cheers and Ettensohn, 2004; Song *et al*., 2012; Stepicheva and Song, 2015). Injection solutions contained 20% sterile glycerol, 2 mg mL^-1^ 10,000 MW FITC lysine charged dextran (ThermoFisher Scientific, Waltham, MA) and 50 ng µL^-1^ of *Renilla* Luciferase (RLuc) constructs. Injections were performed using the Pneumatic pump system (World Precision Instruments, Sarasota, FL) (Stepicheva and Song, 2015; Stepicheva *et al*., 2015). A vertical needle puller PL-10 (Narishige, Tokyo, Japan) was used to pull the injection needles (1 mm glass capillaries with filaments) (Narishige Tokyo, Japan).

The 3’UTR of *Aurora B* was amplified with PCR using sea urchin cDNA and cloned into the ZeroBlunt vector (ThermoFisher Scientific, Waltham, MA) (Table 1 Primers). Positive clones were sequenced (Genewiz Services, South Plainfield, NJ) and subcloned into *RLuc* as described previously (Stepicheva *et al*., 2015). The CPE element was identified bioinformatically and was deleted from the plasmid using site-directed mutagenesis (Lightning QuikChange Mutagenesis, Agilent, Santa Clara, CA). DNA sequencing of these plasmids indicated successful deletion (Genewiz, NJ). The plasmids were digested with EcoRI (ThermoFisher, Scientific, Waltham, MA) and RNA was *in vitro* transcribed using mMESSAGE mMACHINE Sp6 Transcription Kit, ThermoFisher, Scientific, Waltham, MA mRNA was purified using NucleoSpin RNA clean up kit (Macherey-Nagel, Bethlehem, PA), and passed through a Millipore Ultrafree 0.22 μm centrifugal filter (MilliporeSigma, St. Louis, MO) prior to microinjections. RNA constructs were injected at a final concentration of 50 ng µL^-1^.

### Block CPE element with antisense morpholino oligonucleotides (MASO)

To examine if CPE element is important for localization of *Aurora B* transcript to the mitotic spindles, we designed a target protector MASO (TP) blocking the cytoplasmic polyadenylation element (CPE): 5’ AGCTCGAATGATAAAGCTTAC<u>TTTAAAA</u>CA 3’, with CPE sequence underlined (GeneTools, Philomath, OR). Due to the high A-T content of this CPE region, the TP sequence was designed to be a 30-mer to ensure sufficient affinity to the *Aurora B* transcript. For negative controls, we used 5’ CCTCTTACCTCAGTTACAATTTATA 3’, that targets a human *beta-globin* intron mutation purchased from GeneTools (Philomath, OR). We also designed a negative control TP complementary to the 3’UTR of *Aurora B*: 5’ CTCAACATACGTTTTCATACAAAGT 3’ that is upstream of the CPE. Embryos were injected with a final concentration of 5 μM, 50 μM, and 500 μM, and observed at 24 hpf to determine which concentration resulted in 50% mortality. All experiments described were performed using a final concentration of 500 μM.

Embryos were injected with negative control, *Aurora B* TP control and *Aurora B* CPE TPs, then the embryos were assessed for stage of development every hour after fertilization until 6 hpf.

### Drug studies

Physiologic embryos were fertilized and cultured to 16-32 cell stage (∼5 hpf) and were treated with either kinesore 50 µM, ciliobrevin D 100 µM, colchicine 10 mM, cytochalasin D 20 µM, or DMSO at equivalent concentrations in FSW for 30 minutes at 12°C. All drugs are obtained from MilliporeSigma (Burlington, MA) and dissolved in DMSO. The embryos were then fixed immediately and followed by FISH or collected for real time, quantitative PCR (qPCR).

### Real time, quantitative PCR (qPCR)

To examine the relative quantities of transcripts of *Aurora B, Polo kinase, Dynein, Staufen* and *Tubulin α1* after disruption of cytoskeletal dynamics, we used qPCR to examine their transcript levels. Two hundred eggs or embryos were collected immediately prior to treatment with kinesore, ciliobrevin D, colchicine, cytochalasin D or DMSO and immediately after treating for 30 minutes. Total RNA was extracted with NucleoSpin RNA XS kit (Macherey-Nagel, Bethlehem, PA). cDNA was synthesized using the iScript cDNA synthesis kit (BioRad, Hercules, CA). qPCR was performed using 7.5 embryo equivalents for each reaction using Fast SYBR Green Master Mix (ThermoFisher Scientific, Waltham, MA). Reactions were run on the QuantStudio 6 Real-Time PCR cycler system (ThermoFisher Scientific, Waltham, MA), as previously described (Sampilo *et al*., 2018). Threshold cycle (Ct) values were normalized first to *ubiquitin* and shown as fold changes compared with DMSO treated embryos, using the 2^−ΔΔCt^ method (Stepicheva and Song, 2015). Primers were designed using Primer3 (Untergasser *et al*., 2012) (Table 1).

### Image J Analysis

To quantitative analyze the enrichment of transcripts to the mitotic spindle, single plane images of embryos containing blastomeres in anaphase were exported from Zen as jpegs. These images were opened in ImageJ (Schneider, Rasband and Eliceiri, 2012). A region spanning the area between the chromosomes was selected and the mean fluorescence intensity (MFI) was measured, the spindle MFI. An area of the exact same size was selected in the cytoplasm and the MFI of this area was measured, the cytoplasmic MFI. The ratio was then calculated by dividing the spindle MFI by the cytoplasmic MFI.

## Acknowledgements

The authors would like to thank Elizabeth McCulla and other undergraduate students in BISC412 who helped generate the RNA *in situ* probes used in this manuscript. We also thank Nina Faye Sampilo for the *NuMA* FISH images. The authors would also like to thank Dr. Gary Laverty (University of Delaware) for his kind gift of the LLC-PK1 cells. Figures were created with using Biorender.com. We would also like to thank the anonymous reviewers for their insightful comments.

---

**Figure S1:**
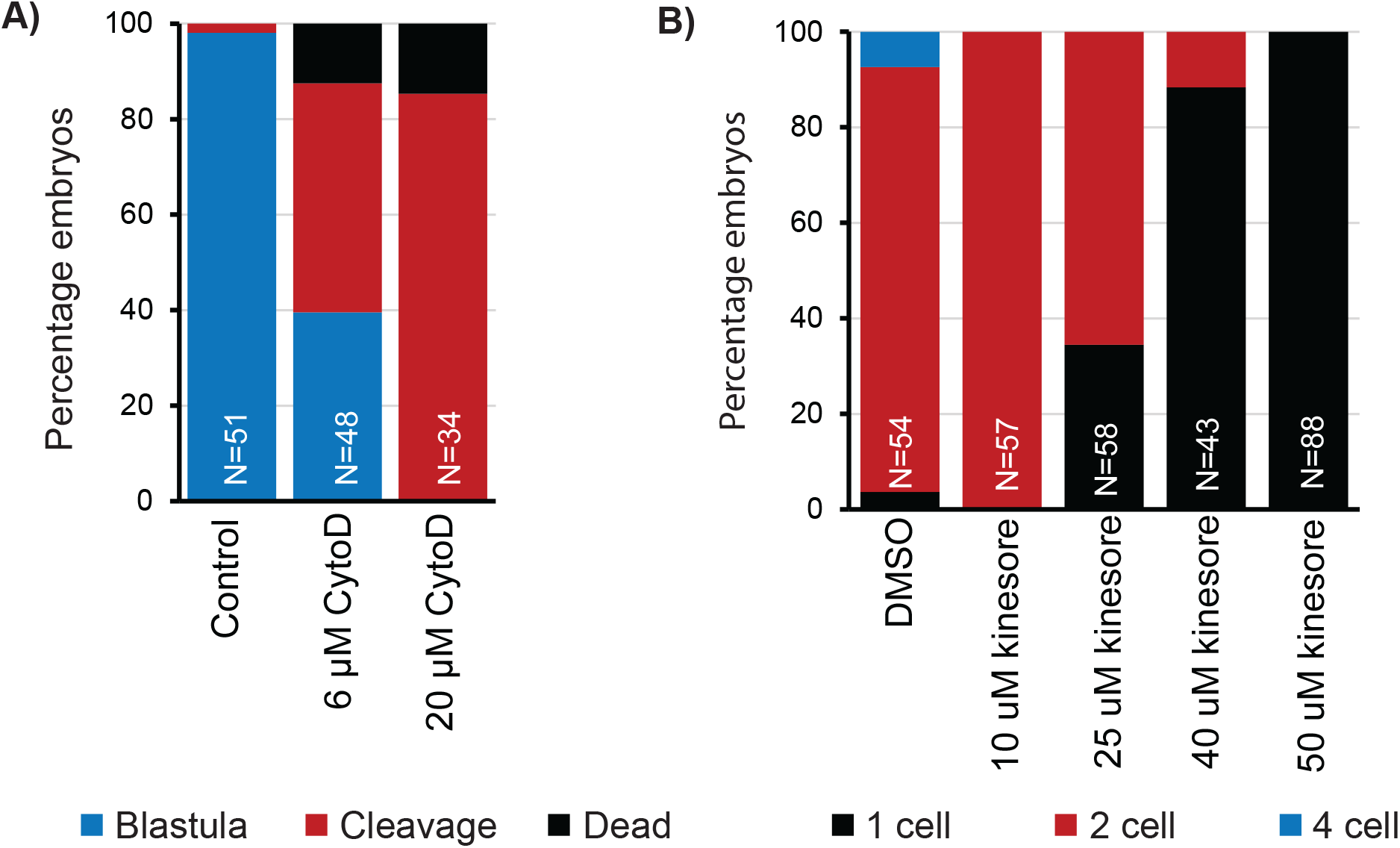
Treatment with cytochalasin D and kinesore result in cleavage stage arrest. (A): Embryos treated with cytochalasin D exhibit cleavage stage arrest. Embryos were treated with the indicated dose of cytochalasin D at early cleavage stage and assessed at 24 hpf to determine survival. (B) Embryos treated with kinesore exhibit cleavage stage arrest. Embryos were treated with the listed dose of kinesore and assessed at 2 hpf to determine survival.

**Figure S2:**
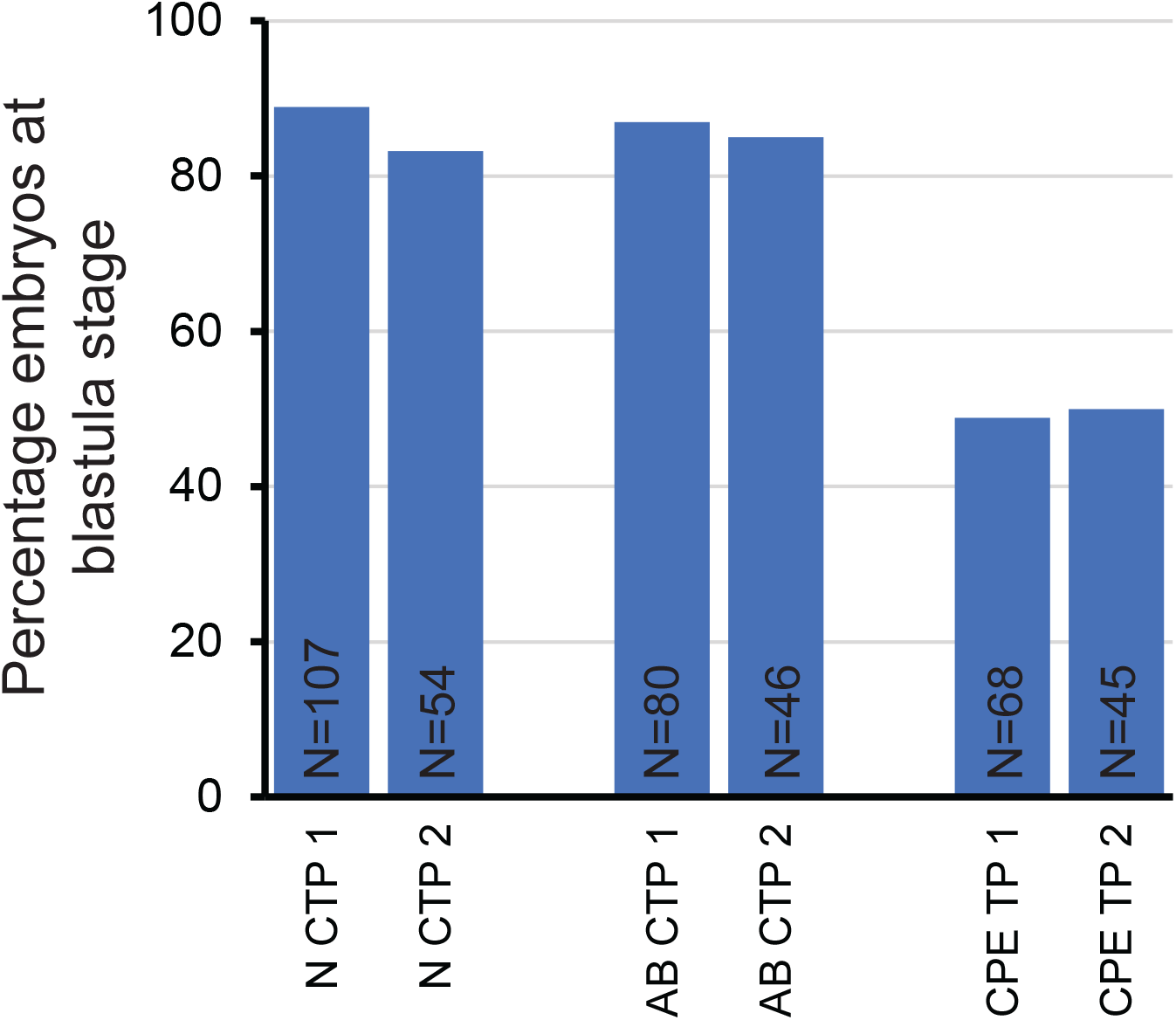
Embryos injected with Aurora B CPE TP exhibit decreased survival to blastula stage. Embryos were injected then allowed to develop to 24 hpf. 2 biological replicates.

